# The Tactile Receptive Fields of Freely Moving *Caenorhabditis elegans* Nematodes

**DOI:** 10.1101/259937

**Authors:** E. A. Mazzochette, A. L. Nekimken, F. Loizeau, J. Whitworth, B. Huynh, M.B. Goodman, B.L. Pruitt

## Abstract

Sensory neurons embedded in skin are responsible for the sense of touch. In humans and other mammals, touch sensation depends on thousands of diverse somatosensory neurons. By contrast, *Caenorhabditis elegans* nematodes have six gentle touch receptor neurons linked to simple behaviors. The classical touch assay uses an eyebrow hair to stimulate freely moving *C. elegans*, evoking evasive behavioral responses. While this assay has led to the discovery of genes required for touch sensation, it does not provide control over stimulus strength or position. Here, we present an integrated system for performing automated, quantitative touch assays that circumvents these limitations and incorporates automated measurements of behavioral responses. *Highly Automated Worm Kicker (HAWK)* unites microfabricated silicon force sensors and video analysis with real-time force and position control. Using this system, we stimulated animals along the anterior-posterior axis and compared responses in wild-type and *spc-1(dn)* transgenic animals, which have a touch defect due to expression of a dominant-negative *α* spectrin protein fragment. As expected from prior studies, delivering large stimuli anterior to the mid-point of the body evoked a reversal, but such a stimulus applied posterior to the mid-point evoked a speed-up. The probability of evoking a response of either kind depended on stimulus strength and location; once initiated, the magnitude and quality of both reversal and speed-up behavioral responses were uncorrelated with stimulus location, strength, or the absence or presence of the *spc-1(dn)* transgene. Wild-type animals failed to respond when the stimulus was applied near the mid-point. These results establish that stimulus strength and location govern the activation of a stereotyped motor program and that the *C. elegans* body surface consists of two receptive fields separated by a gap.

## Introduction

Sensory mechanisms underlie many of the interactions among living things and with the physical world. Touch and pain sensation depend on sensory neurons embedded within and distributed throughout the skin. The size and shape of tactile receptive fields and how sensory neurons tile the skin surface are critical factors governing the spatial resolution and sensitivity of the somatosensory system ^1-3^.

The six touch receptor neurons (TRNs) of the nematode *Caenorhabditis elegans*, ALML, ALMR, AVM, PLML, PLMR, and PVM, comprise a set of mechanosensory neurons that tile the body surface and link gentle touch sensation to behavior ^4,5^. The TRNs are needed for touch-evoked avoidance behaviors; a simple circuit links TRN activation to interneurons and motor neurons ^6,7^. From optogenetic dissection studies, TRN activation is known to be sufficient to activate such behaviors: stimulation of the three anterior neurons (ALML, ALMR, and AVM) causes forward-moving animals to reverse direction, while stimulation of two posterior neurons (PLML and PLMR) results in a speed-up ^8^. A third posterior neuron, PVM, is activated via mechanical stimulation ^9-11^ but is not required for behavioral responses to gentle touch ^6,7^. Thus, five of the six TRNs are necessary and sufficient to produce touch-evoked avoidance behaviors in adult hermaphrodites.

Classical touch assays use a tool consisting of an eyebrow hair mounted on a toothpick to stimulate (touch) an animal, after which the observer visually scores whether the animal reverses direction ^12^. This assay has been used to identify dozens of genes required for touch sensation ^5^ and has a number of advantages. For example, it is a non-invasive measurement of freely moving and intact animals and the experimental setup is straightforward. This assay also has several limitations. First, the strength of the touch stimulus is neither known nor controlled. Rather, stimulus strength depends on manual control of the touch tool and the mechanical characteristics of the eyebrow hair ^13^. Second, the position of the stimulus on the body surface is unknown and its spatial precision is limited. Third, posterior touch is expected to accelerate forward movement (speed-up), which is not easy to score quantitatively by observation. This situation limits our knowledge of the relationship between posterior TRN stimulation and speed-ups as well as the precise spatial distribution of *C. elegans’* touch sensation. Using a microfluidic device, McClanahan, et al. recently demonstrated a microfluidics-based behavioral touch assay which locally compressed the worms using a pressurized membrane at the top of the channels ^14^.This system was deployed to score worm behaviors and quantify differences in the receptive fields of wild-type animals and those with defective posterior TRNs. This system provided detection stimulus location but no readout of the stimuli force or deflection into the animal’s body.

Although the general relationships among TRN tiling, TRN tactile receptive fields, and avoidance behaviors are well-known, neither the precise boundaries of the worm’s tactile receptive fields nor the logic that links TRN activation to reversal or speed-up are known. To generate the data needed to fill this critical gap, we developed the *Highly Automated Worm Kicker* (HAWK). HAWK tracks freely moving worms, delivers controlled mechanical stimuli to specific positions along the anterior-posterior axis, and automatically classifies behavioral response types (pause, reversal, speed-up, and null responses). Central to the function of HAWK are two key, custom innovations: 1) a custom-force sensing cantilever that acts as the stimulator ^15-17^ and 2) an optical system to track and target the animal. The detailed design, fabrication and signal conditioning of these silicon, piezeoresistive cantilevers were previously reported ^18-20^ We present the design of the optical system here. We used HAWK to systematically vary both stimulus position and strength, mapping tactile receptive fields and analyzing the relationship between these stimulus parameters and the nature and intensity of motor responses.

As anticipated, stimuli delivered anterior to the mid-point of the worm body evoked reversals, while those delivered posterior to the mid-point triggered speed-ups. Unexpectedly, the anterior and posterior receptive fields were separated by a gap in which even the strongest mechanical stimuli failed to reliably evoke a reversal, a speed-up, or a pause. Touch sensitivity varied within the anterior and posterior receptive fields: positions distal to receptive field boundaries were more sensitive to mechanical stimulation. Additionally, we found that stronger stimuli more reliably evoked motor responses but that the intensity of such motor responses was not correlated to stimulus strength. Thus, signaling in the TRN-motor circuit reflects execution of a stereotyped motor program. This finding suggests that the TRNs function as a probabilistic switch between two states: the neuron is either signaling to execute the motor program or is not. Our results suggest that the probability of evoking a given response type depends on stimulus strength and location, but once executed, responses appear to consist of stereotyped, invariant motor programs.

## Methods

### Strains

The following *C. elegans* strains were used: wild-type (N2), CB61 *dpy-5(e61)* I ^21^, CB678 *lon-2(e678)* X ^22^, and GN539 *pgIs10 [Pmec-17::spc-1[1-170]::mCherry]* ^23^. We refer to GN539 as TRN::SPC-1(dn) for convenience and to emphasize that this transgenic line expresses a dominant-negative fragment of the SPC-1 α spectrin exclusively in the TRNs. Note that these strains yield nearly pure populations of hermaphrodites, if males arise in the populations they are not used in the studies. Animals were cultured on standard NGM plates covered with a lawn of OP50-1 bacteria at 20 °C ^24^. The worms were synchronized 48 h before experiments, to ensure that all animals were late L4 larvae or young adults, i.e., when AVM and PVM are expected to be synaptically connected.

### Sample preparation

For each sample, we cleaned a 75 × 50 mm glass slide and a 50 × 24 mm, 0.13-0.17 mm thick cover slip via sonication in 2% Hellmanex solution for 5 min, rinsed them in deionized water, and dried them. The sample pad consisted of 180 μL of melted 2% NGM agar pipetted onto a glass slide between two strips of tape; the pad was flattened to a consistent thickness by placing a cleaned cover slip across the tape. To transfer animals with minimal bacterial food, we washed well-fed animals from a growth plate into a microcentrifuge tube in sterile water, allowed the animals to settle to the bottom of the tube, transferred 20 μL of a concentrated animal suspension to a piece of filter paper (Whatman, Grade 1, 5.5 cm diameter), and inverted the filter paper onto an unseeded NGM plate. Using this procedure, we typically transferred 15-20 animals per 20 μL. After 10-15 min of recovery, single animals were transferred with a platinum pick to a freshly prepared experimental sample pad.

### Automated touch assays with HAWK

We prepared samples of worms as described above and placed them on the stage of our custom system, HAWK. In a typical experiment, we collected 3-4 bright-field images of each animal and then made three 30 s recordings of behaviour with no mechanical stimulus. To set up the assay, we selected the target location on the body and the force profile and started tracking. When the animal was moving forward, we initiated the force application. In this way, animals were in the same locomotion state prior to stimulation, independent of the target stimulus position. We used a the real time, closed loop force clamp system to apply a step force profile vertically onto the animal, and maintain the desired force for 100 or 150 ms ^16^. The cantilever was then positioned to maintain contact with zero force for a 2-s dwell interval because pull-off of the force probe from a wetted surface generates significant force which itself can elicit a behavioral response ^25^. We paused stage movement during stimulus application to prevent electrical coupling between the stage motors and the force probe-sensing signal. HAWK recorded behavior for 10 s after the stimulus. We applied as many as two stimuli per minute, until we had collected data from 12-13 trials. We then switched to another target/force combination. We continued with this method for 1-1.5 h per worm.

### HAWK system design

We developed a fully automated, quantitative touch assay system, HAWK, by integrating two technologies: 1) real-time tracking and 2) mechanical stimulus control. For tracking and targeting, we built on systems reported by Leifer et al. ^8^ and Stirman et al. ^26^, who implemented real-time tracking for delivering optical stimuli to user-defined body segments (Figure 1). For mechanical stimulus delivery, we relied on our previous system consisting of a custom, self-sensing, cantilever force probe ^18-20^ integrated into a closed loop control system ^16^. HAWK introduces the ability to automatically measure and classify behavioral responses. Thus, HAWK implements two features absent from prior systems ^15,25^ for the study of mechanosensation: 1) the delivery of controlled forces to specific body positions in freely moving worms and 2) automated analysis of behavioral responses.

**Figure 1.**
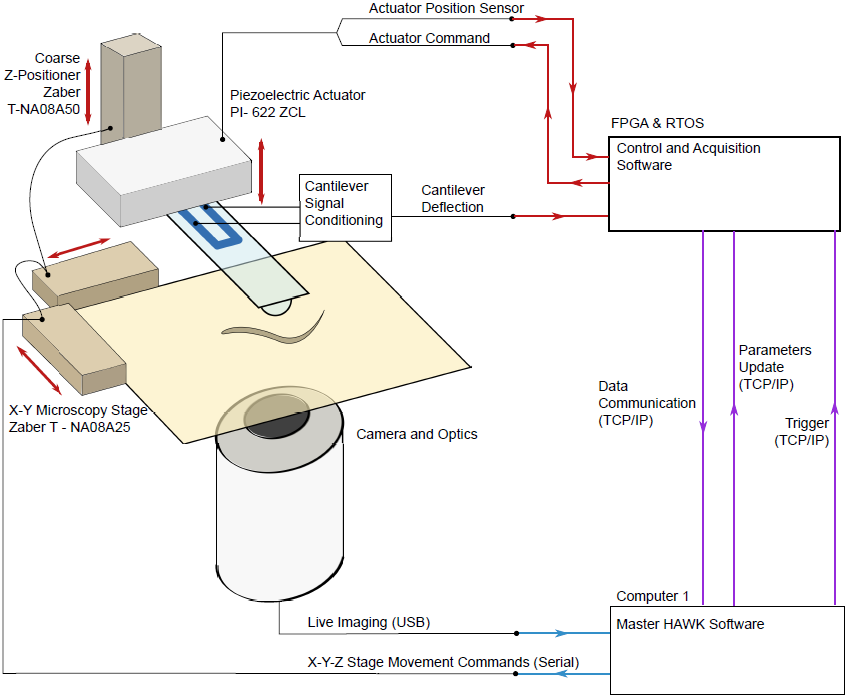
HAWK system overview. A PC computer runs the custom HAWK software to manage the tracking, stimulus set up and trigger, data acquisition, and user interface. The software controls the camera and stage hardware. The oblique light source and inverted configuration of HAWK images worms, but not the force probe or the background. The camera captures images of the freely moving animal, which are processed by the HAWK image processing module. The HAWK software then translates the microscopy stage to position the animal underneath the force probe, enabling targeting. To apply a controlled force, the RTOS/FPGA uses the sensor signals from the force probe displacement and the actuator position to adjust the piezoelectric actuator position.

### HAWK hardware

HAWK imaging hardware consists of two SLR camera lenses (Nikon) mounted on a CCD microscope camera (The Imaging Source, DMK 31AU03), an automated microscopy stage (ASR100B120B-T3, Zaber), and a cantilever mounted above the stage (Figure 1). We inverted the upper lens for magnification, focused the second lens at infinity, and positioned the lenses and camera below the sample stage. An array of red LEDs (Super Bright LEDS) illuminated samples from the side, creating an imaging system akin to dark field microscopy. Since the cantilever is not in the light path, the lenses capture light scattered by the animal and leave the background and cantilever dark. This illumination method made real-time tracking compatible with force delivery via our feedback-controlled force-sensing cantilever.

HAWK force delivery hardware consists of a custom-designed silicon cantilever fabricated with an integrated piezoresistor at the base of the cantilever. In this way, resistance changes are proportional to the displacement of the cantilever tip ^27^. These signals are related to the force applied using the spring constant of the cantilever ^18^. We mounted cantilevers on a custom printed circuit board for signal conditioning. As illustrated in Figure 1, the circuit board-cantilever package was mounted on a piezoelectric actuator (Physik Instrumente Inc. model 622.ZCL) for precision feedback-control of the cantilever vertical position and concomitant force. Gross cantilever positioning was controlled with a motorized vertical stage (462-Z-M, Newport, T-NA08A5, Zaber) in concert with manual XY linear stages (SM-50, M-433, Newport).

We measured the resistance change in the piezoresistor embedded in the cantilever using a Wheatstone bridge configuration in series with an instrumentation amplifier (INA103, Texas Instruments). The resulting voltage signal was proportional to the cantilever tip displacement and thus applied force. To define a predictable contact surface between the cantilever and the animals, we glued a 10-μm glass bead to the tip of each cantilever used in this study (Figure 2A). Because the cantilever contacts animals from above and because *C. elegans* nematodes crawl on their sides, an ALM (or PLM) neuron was below the glass bead in anterior (or posterior) touch assays. We coated cantilevers and the attached glass beads with ∼400 μm of parylene-N (Specialty Coating Systems) for electrical passivation. We calibrated each cantilever via Laser Doppler Vibrometry to determine its resonant frequency (f_0_, Hz) and spring constant (k, N/m) ^28,29^.

**Figure 2.**
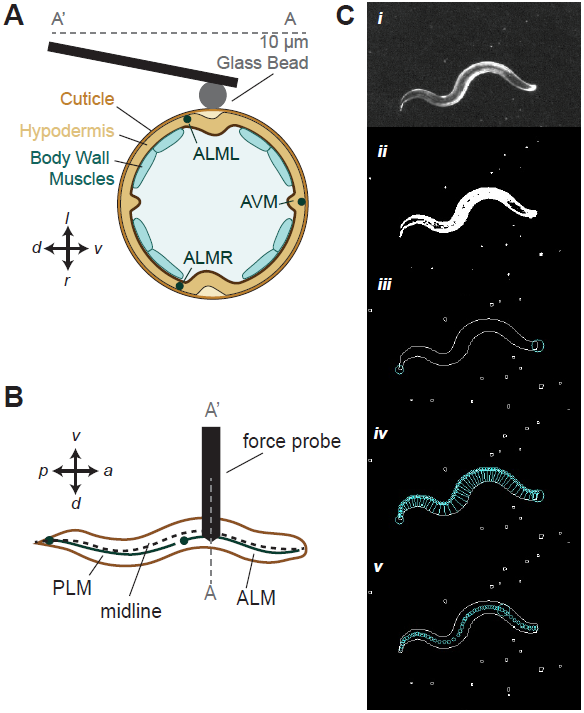
Force probe actuates vertically to apply controlled force profiles along the animal’s midline. A) Cross-section schematic of experiment showing contact between animal and the glass bead attached to the force probe, which is positioned near the lateral midline and moves vertically to indent freely moving animals. B) Top view schematic of the touch assay that targets the ALM or PLM neurons along the midline. C) Image processing pipeline for detecting the target on the animal. Details of process steps are described in the Methods section.

We determined the displacement sensitivity (V/m) of each cantilever by pressing the tip of the cantilever against a glass slide and calculating the slope of the relationship between actuator displacement and cantilever sensor voltage signal. The actuator drive signal, the actuator displacement sensor signal, and the cantilever deflection signal were integrated into a proportional-integral-derivative feedback loop programmed in the real-time operating system of a field programmable gate array (RTOS/FPGA, model CompactRIO, National Instruments). We tuned the PID control parameters for specific force profiles using a calibration module coded in the HAWK software.

### HAWK software

The HAWK software is based on a Windows Form Application to control experiments (tracking, targeting, behavioral analysis) and to record experimental data in YAML format. The software was designed to run on a Windows 7 PC (Dell Precision T1700 Workstation) ^30^. It includes a graphical user interface that displays a live image from the digital camera, stage position controls, coarse vertical positioning of the force probe, and experiment controls including a setup dialog box. Three modules work in parallel while the application runs: 1) tracking, 2) force clamp control, and 3) data management. The tracking module captures images, performs image processing, and controls x-y stage position in real time. The force clamp module manages communication with the RTOS/FPGA for the force clamp ^16^. The data management module records image processing and stage movement data generated in the HAWK software as well as experimental parameters and stimulus profile data from the force clamp module; it saves data to the hard disk in real time. Table 1 summarizes the average time needed to perform each of the operations handled by HAWK software.

**Table 1.** HAWK Processing Time. Average and standard deviation (SD) of major processing time steps in tracking loop across 450 experiments consisting of 4,229,026 tracking loops in 2,295 trials.

| Processing Step | Average (ms) | Standard Deviation (ms) |
| --- | --- | --- |
| Image Acquisition | 19.556 | 1.034 |
| Find Worm | 21.307 | 2.203 |
| Move Stage Command | 5.492 | 0.021 |
| Wait For Stage | 46.417 | 2.983 |
| Complete Tracking Loop | 107.007 | 4.167 |

We used the HAWK tracking and targeting software modules to apply controlled forces to a freely moving worm at a user-specified location along the animal’s anterior-posterior axis, or midline. We achieved this precision by tracking the midline of a freely moving animal using a strategy adapted from Leifer, et al. ^8^ to extract the midline of the animal and to compute the user-defined target location on the midline (Figure 2B). The force probe was positioned over the target position by translating the microscope stage in real time. We also recorded the position of the midline for post-hoc analysis of behavior response.

The processing pipeline for extracting the midline of moving animals is applied in real-time and involves the following steps (see also Movie 1 and Figure 2C, from top to bottom). 1) Use Otsu’s method to compute a threshold for segmenting the image ^31^. 2) Define the largest foreground object as the worm.3) Find the contour to obtain a list of pixels along the outside of the body. 4) Determine which end is the head and which is the tail by calculating sharpness at each point on the contour, S

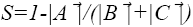 where 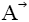 is the vector between two points that are each 10 points along the contour on either side of the current point; 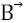 and 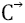 are vectors between the current point and the two points that are each 10 points along the contour on either side of the current point. The tail is the sharpest point while the head is the sharpest point in the one quarter of the points along the contour opposite the tail. 5) Search for pairs of points on opposite sides of the body contour with minimum distance between them to create segments across the body. Start at the head and move along one side of the contour until reaching the tail.6) Define the midline as the segment midpoints. 7) Calculate the target as a percentage of the total distance of the skeleton, the sum of the distances between the skeleton points. The final target pixel location is linearly interpolated along the segment between two skeleton points in front of and behind the target. Movie 1 is an example of how HAWK integrates tracking and spatially precise mechanical stimulus delivery.

### HAWK-based measurements of body morphology

We calculated the body length of the animals from the sum of the distances between points along the skeleton. The body width was taken from the length of the segment closest to the midpoint of the skeleton, or 50% of the animal’s body length.

### Automated analysis of mechanically evoked behavior

To detect responses to mechanical stimulation, we devised methods for classifying responses as a speed-up (forward acceleration), a pause, a reversal, or a null response; we also established methods to measure the intensity of these responses. We distinguished forward from backward locomotion by taking advantage of the fact that a curvature wave propagates head-to-tail during forward *C. elegans* movement and tail-to-head during backward movement ^32^. Thus, the velocity of the animal, V(t), is the velocity of curvature propagation along the midline, as previously described ^8,30^. The direction of movement is directly related to the sign, positive or negative, of the velocity.

To determine the velocity of the bending wave as a function of time, *V*(t), we compared the curvature in the current frame, *n*, with that the previous frame, *n-1*, to find the phase shift in the body curvature. We used a least-squares fitting algorithm to find the phase shift that minimized the root-mean-squared error between the contours in frame *n* and *n-1*. The phase shift is related to the velocity by the number of points sampled for the curvature and the animal’s body length ^8,30^. We smoothed the bending wave velocity trace with a one-dimensional Gaussian kernel with sigma of 1.5.

Next, because we are primarily interested in stimulus-induced changes in *V(t)*, we defined a normalized measurement of velocity as a function of time for each trial (Eqn 1):

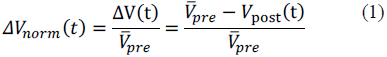

where 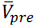 is the average velocity 1.5 s prior to stimulation and v_post_(t) is the velocity as a function of time of the animal after the stimulus is applied. Because stimuli are applied only during forward movement, 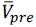 always has a positive value. We relied on v_post_ (t) to classify responses and *ΔV/V* to measure response intensity (Eqn 2):

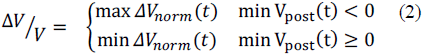

It follows somewhat counter intuitively from the initial condition of forward movement and Equation 1 that if the animal speeds up, then *ΔV/V* has a negative value and that if the animal reverses direction, *ΔV/V* is positive.

### Criteria for data inclusion and exclusion

Several criteria were applied as quality-control measures for data inclusion. The first criterion was related to the animal’s behavioral state: we required that each worm be moving forward at an average speed of at least 50 μm/s during the 1.5 s before the stimulus application. A second criterion was related to stimulus targeting accuracy. In particular, we required that the cantilever’s tip contacted the animal within 10% of the target position as a function of length and within the middle 50% of the animal’s width. The third set of criteria is related to the tracking algorithm, which occasionally failed to accurately identify the animal’s midline. We used the width, length, and continuity of the skeleton to flag tracking failures on a frame-by-frame basis. Specifically, frames in which the skeleton was more than 125 μm wide or <85% of the average length were flagged as failures because these dimensions are unrealistic outcomes for the animals we studied. Frames in which the skeleton intersected with itself were also flagged as failures in skeleton integrity. We excluded trials in which tracking failed in more than 6/22 or 27% of the frames in a given trial. In total, 89.4% of trials succeeded in meeting all three set of criteria and were included in our data set.

There were two classes of reversals identified as false positives by human inspection. The first class was triggered by removal of the cantilever after the zero-force dwell interval (2 s). In these cases, the response was evoked by breaking adhesion between the cantilever and the animal, rather than by the stimulus delivered earlier in the trace. The second class consisted of false reversals due to errors introduced into the velocity calculation during a large translation of the animal between two frames at the end of the stimulus: since we could not actively track moving animals during stimulus delivery, therefore we occasionally needed a large stage translation to bring the animal back under the cantilever after the stimulus. However, we could not capture images during this time due to motion blur. Fewer than 20% of the trials required a manual adjustment for both classes of false positive reversals.

### Comparing automated behavioral classification to human observation

Five human observers reviewed videos of responses to 188 distinct stimuli encompassing five forces between 100 nN and 10,000 nN and three body targets: 25%, 35%, and 45% of the distance from the head to the tail. Three observers were novices having no prior experience with *C. elegans* behavioral assays and two were experts with prior experience performing and analysing *C. elegans* behavioral assays. The videos were coded such that although the observers were unaware of the magnitude of the stimulus applied, they could visually observe the force probe as it came in contact with the animal. Each person received a scoring template with the order of the trials randomized and was asked to classify the type of response from the animal as a reversal, a pause, a speed-up, or a null response.

### Statistical Analysis

To measure the agreement rate of behavior response scoring we used the Cohen’s kappa coefficient, which measures agreement rate between two scorers (human-human pairs and human-computer pairs) by accounting for the probability of the two raters randomly agreeing ^33,34^. We calculated the mean kappa coefficient and corresponding standard error of the mean among human-human pairs and then human-computer pairs.

To perform the best fit analysis of the behavior response probability, we fit the measured response rate according to, *f(x) = a(1-e*^*-bx*^). The fit was weighted by the inverse of the variance around the animal’s mean response rate to the corresponding force in the same method as Petzold, et al ^25^.

To compare each of the behavioral response metrics, *ΔV/V* and maximum acceleration we performed an ANOVA analysis using the Matlab ™ method “anovan,” with a type III sum of squares. For reversals, we have two factors: 1) stimulus strength, and 2) target location, so we used a two-way ANOVA analysis. When characterizing the metrics for speed-ups, we used one target, 75%, so we used a one-way ANOVA analysis with stimulus strength as the single varying factor. Finally, when comparing the metrics for reversals between wild-type animals and *spc-1(dn)* transgenic animals, we used a two-way ANOVA analysis with the factors of stimulus strength and genotype. P-values less than 0.01 were considered significant.

## Results

### Force delivery and targeting precision

Classical touch assays rely on a manually controlled eyebrow hair to deliver mechanical stimuli ^12^. The forces delivered by human experimenters wielding an eyebrow hair are highly variable, but generally exceed 10,000 nN ^13^. By contrast, HAWK delivers repeatable and controlled stimuli, with forces as low as 50 nN and rise times in the range of 20 ms (Figure 3A). HAWK is also designed to deliver mechanical loads to user-defined locations along the anterior-posterior body axis. In this study, we exploited these features to map the tactile receptive fields of wild-type *C. elegans* hermaphrodites and to determine whether sensitivity to mechanical loads is uniform or variable within a given receptive field.

**Figure 3.**
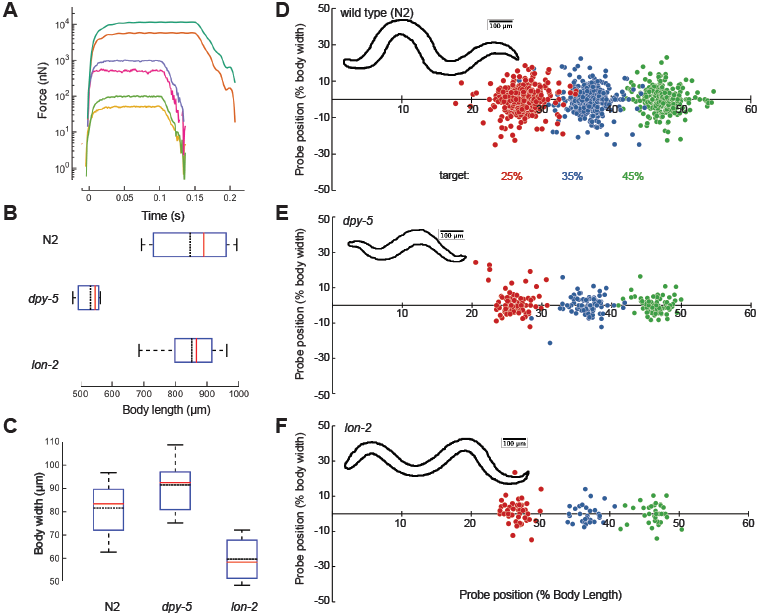
Precision and robustness of force applied and position by HAWK. A) Single, representative force profiles applied to freely moving worms spanning 50 nN to 10,000 nN. For larger forces, we extended the pulse length to 150 ms to account for the longer rise time required to reach full stimulus strength. B) Body length measurements of wild type (N2), *dpy-5*, and *lon-2* animals determined from the length of the midline. C) Body width measurements made at 50% of the body length for all three genotypes. Box plots show median (red line), the 25% and 75% percentiles and whiskers at 10% and 90% percentiles; the black dotted line is mean. *N =* 66, 15, and 14 measurements from 33 wild type, 4 *dpy-5*, and 5 *lon-2* animals. D-F) Stimulus location at stimulus onset according percent of the body length and width from the midline for wild type across all trials from 33 wild type, 4 *dpy-5*, and 5 *lon-2* animals (8-12 trials per location per animal) (D), *dpy-5* (E), and *lon-2* (F). The sign of the y-coordinate is assigned randomly. Circle color indicates the desired target position (in %): 25 (red), 35 (blue), 45 (green). Insets show representative body contours for each genotype.

To assess the precision of the stimulus-targeting feature, we measured the position of the cantilever tip relative to the midline skeleton and its normalized position along the anterior-posterior axis for stimuli directed at 25%, 35%, and 45% of body length (the tip of the nose is 0% and the tail is 100%). To assess the robustness of the stimulus-targeting feature, we compared precision for wild-type animals with the precision achieved for *dpy-5* and *lon-2* mutants, whose body shapes differ from that of wild-type worms ^21,22^. *dpy-5* animals are both shorter and fatter than wild-type animals; *lon-2* animals are thinner and longer (Figure 3B, 3C). For each target location and genotype, we applied 8-12 stimuli per animal. For wild-type animals, the vast majority (95%) of stimuli were applied within 10% of the body length (∼100 μm) and 25% of the body width (∼20 μm) of the desired target. Note that 89% of the stimuli were within 5% body length or ∼50 μm (Figure 3D). Deviations from the target position were biased posteriorly due to the forward movement of the animal during the time that stage motion was halted. This pause in stage motion was needed to minimize electrical interference between the stage motors and the cantilever displacement signal. Nevertheless, targeting accuracy was similar in wild-type, *dpy-5*, and *lon-2* animals (Figure 3E, 3F), indicating that HAWK delivers defined mechanical loads to freely moving animals within 10% of a desired target in a manner that is robust to substantial changes in body shape.

### Automated classification of behavioral responses

*C. elegans* propel themselves via alternating dorsal-ventral muscle contractions that produce propagating waves. We determined the animal’s propagating wave velocity (*V*) by calculating the phase shift between the midline curvature between frames (Methods). Four types of behavioral responses were classified from *V* traces in the 2-s post-stimulus period (Methods): a speed-up, a pause, a reversal, or a null response (Figure 4A). A trial was classified as a reversal if the velocity crossed zero during the post-stimulus period. If no reversal was detected, then the algorithm analyzed the velocity trace for a pause event, defined as any trial in which speed fell below 30% of the pre-stimulus speed. If neither a reversal nor a pause was detected, then we tested for a speed-up by calculating the forward acceleration profile of the animal using the derivative of the velocity, *ΔV(t)/dt*. If the maximum acceleration in the post-stimulus period was >400 μm/s^2^, then the response was classified as a speed-up. We determined the 400 μm/s^2^ threshold empirically. First, we measured the acceleration of ∼30 animals during 30-s forward runs. For each animal, we calculated the mean and standard deviation of the acceleration measurements during the run. The average of the standard deviations measured for each of 30 animals was 200 μm/s^2^. We set the threshold to two times this average. If the velocity profile was stable and inconsistent with a reversal, a pause, or a speed-up, then we recorded the trial as a null response.

**Figure 4.**
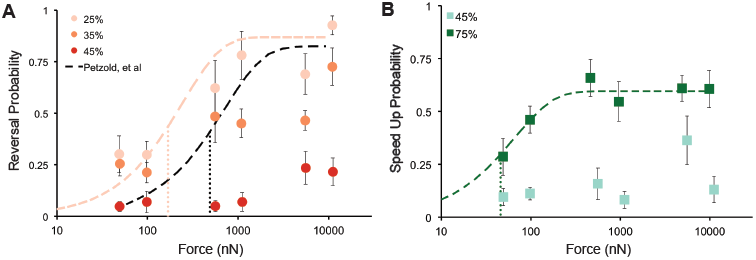
*C. elegans* adults execute a speed-up, pause, reversal or null response as a function of stimulus strength and location. A) Velocity *vs*. time for four typical behavior responses (speed-up, pause, reversal, response failure). *t=0* is the stimulus start time. The blue line indicates the pre-stimulus average velocity. B) Distribution of *Δ*V/V *vs.* stimulus strength, pooled across all stimulus location. Width of plot represents the relative distribution of *Δ*V/V. C) Running average of *Δ*V/V vs. stimulus location across all trials for high stimulus strengths. Each point corresponds to the average *Δ*V/V from all trials with a stimulus location inside a window size of 7.5% of body length around the location specified on the x-axis.

To quantify the change in behavior before and after the stimulus as a function of stimulus location, we calculated the normalized velocity change, *ΔV/V*, for each trial (Methods). Following from Equation 1 and because the worms were always moving forward prior to stimulation (Methods), there are four ranges of *ΔV/V* that correspond to behavioral responses:

*ΔV/V* < 0 corresponds to fast forward movement, suggesting a *speed-up,*

*ΔV/V* > 1 corresponds to backward movement, suggesting a *reversal*,

0 < *ΔV/V* < 1 corresponds to steady or slow forward movement, suggesting a *pause,* and

*ΔV/V* ∼ 0 corresponds to a *null response.*

### Tactile sensitivity varies within and between receptive fields

We used HAWK to test the hypothesis that behavioral responses vary with both stimulus strength and location. Given that the anterior and posterior TRNs detect forces in the nano-Newton to micro-Newton range ^35,36^, we delivered force pulses with values between 50 nN and 10,000 nN. We measured force-response curves at three anterior body positions (25%, 35%, and 45% of the distance from the head), and two posterior positions (55% and 75% of the distance from the head). Force profiles consisted of a 100-ms pulse (Figure 3A) followed by a 2-s dwell interval at zero contact force. For larger forces, we extended the pulse length to 150 ms to account for the time required to reach full stimulus strength.

Figure 4B shows the distributions of *ΔV/V* for each stimulus strength across all positions. As expected, the distribution of *ΔV/V* was multi-modal. The first mode of *ΔV/V* was centered on zero (pauses, null responses) and extended into negative values (speed-ups), while the second mode consisted of values greater than one (reversals). Note that due to the nature of *ΔV/V* calculation, the vertical distributions did not linearly reflect the vigor of speed-up and reversal. At low stimulus strengths, the first mode of *ΔV/V* was strongly concentrated around *ΔV/V* = 0 and the second mode was small, indicating a low probability of response to smaller forces. As stimulus strength increased, we observed an increase in the spread of both distribution modes. Dispersion of the first mode correlated with an increase in speed-ups evoked by posterior TRN activation, while dispersion of the second mode correlated with an increase in reversals evoked by anterior TRN activation.

Figure 4C shows *ΔV/V* versus stimulus location for all stimulus strengths greater than the force required to evoke a response in 50% of trials, *F*_1/2_ = 490 nN, in wild-type animals ^25^. By analyzing trials with high stimulus strengths, we rejected trials that yielded a null response, enabling us to assess how behavioral responses varied with stimulus location. On the anterior portion of the body, *ΔV/V* >1 corresponded to a high probability of evoking a reversal. On the posterior portion of the body, at ∼75% of the body length, we observed a high probability of evoking a speed-up, suggested by the decrease in *ΔV/V*. Interestingly, between the 45% and 55% stimulus locations, *ΔV/V* was close to zero, despite the high stimulus strength. This region of low touch sensitivity at the center of the animal’s body suggests the presence of a gap between the receptive fields of the anterior and posterior TRNs. Further, the continuous decay of *ΔV/V* between 30% and 45% of the body length indicates a location-dependent probability of evoking a reversal response that suggests reversal probability declines for touches delivered closer to the ALM cell body.

### Stimulus location governs probability of reversal, speed-up, or null response

We classified behavioral response types by applying our scoring algorithm to the wave velocity profiles of wild-type animals and compared its performance to manually-classified behavioral responses (Methods). We compared the data from human and software-based scorers by computing the Cohen’s kappa coefficient for each human-human pair and each human-software pair ^33,34^. On average, the ten pairs of human raters had kappa values of 0.44±0.06 (mean ± sem) and the five pairs of human-software raters had kappa values of 0.28 ± 0.04 (mean ± sem) indicating moderate agreement among human raters and fair agreement between human raters and the automated response detection system. We speculate that the reduced agreement for human-software pairs reflects the difficulty that human raters, but not the software, face in detecting speed-up events. There was high agreement among both human-human pairs and human-software pairs in identifying reversals.

We examined reversals and speed-ups as a function of force, aggregated by target position (Figure 5). Pauses were not analyzed in depth, since this response type constituted <4% of the responses at any of the targets. Figure 5A shows the probability of a reversal according to force and position. Wild-type worms showed high sensitivity at the 25% target. Even at forces as low as 50 nN, the probability of evoking a reversal was better than one out of four trials (0.28). Reversal probability saturated near 0.8 for forces > 1,000 nN. Stimuli delivered at 35% of body length, posterior to the 25% target, appeared to saturate at a lower maximal reversal probability, suggesting a non-uniform sensitivity to touch within a given receptive field. This observation is consistent with the *ΔV/V* decay we found between 35% and 45% of the body length (Figure 4C). Stimuli delivered near the animal’s mid-point (45% target position) rarely evoked a reversal: probabilities were never higher than 0.24, even for stimuli that delivered 10,000 nN of force—a value that is ten times greater than that which saturates reversal probability at the 25% target.

**Figure 5.**
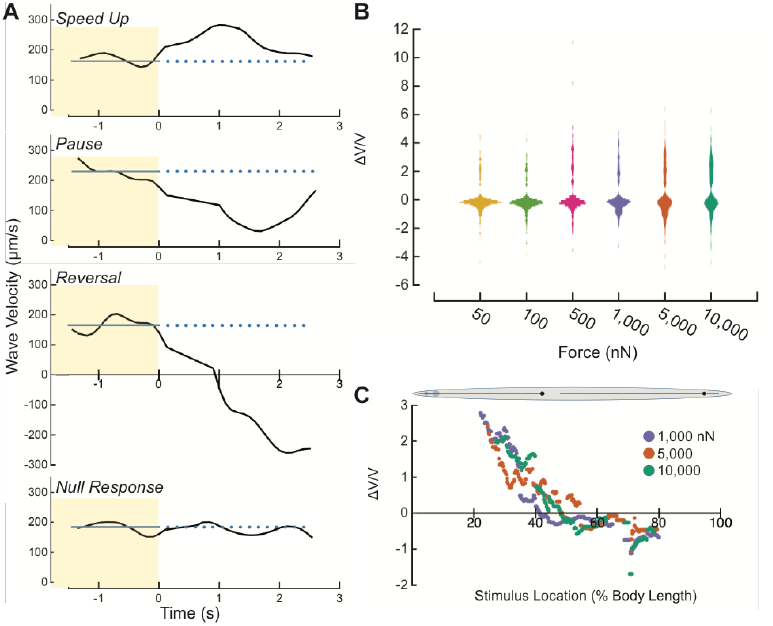
Response type and probability depend on stimulus position. A) Reversal probability *vs.* force for anterior touch. Black dotted line is the fit to the data representing reversal probability from Petzold, et al. ^25^. Pink dotted line was fit to the values recorded at the 25% target. B) Speed-up probability *vs.* force for posterior targets. Green dotted line was fit to the values recorded at the 75% target. Points are the mean response rate ± s.e.m of at least 6 animals. Each animal was tested during at least 4 trials. All fits were performed according to, *f(x) = a(1-e*^*-bx*^), weighted by the inverse of the variance to the means ^25^. Finely dotted vertical lines indicate *F*_*1/2*_ for the respective fit. The values were for 25% and 75% were 168 (95% Confidence Interval: 5.2 – 936) and 46 (95% Confidence Interval: 32 – 63) nN, respectively. We calculated the error in *F*_*1/2*_ by analytically solving for the force that gave the 95% confidence intervals around the half maximal response probability, as determined by the fit error.

Petzold, *et al.* previously measured reversal probability as a function of stimulus force, but stimulus position was controlled manually via a joystick-controlled x-y stage ^25^. While this approach allowed experimenters to control stimulus strength, positioning was less precise and their acceptable range spanned the entire body surface between the 25% and 50% body length. To compare the two datasets, we plotted the fit derived by Petzold, *et al*. with the present data (Figure 5A). The fit reported previously falls in between our 25% and 35% data, consistent with the idea that the prior work measured an average of the response probabilities expected for stimulation of these two body positions (Figure 5A). Using the same method as Petzold, *et al*., we fit an exponential function weighted by the inverse of the variance to our means at the 25% target. We also calculated *F*_*1/2*_ = 168 nN (95% CI: 5.2 – 936 nN) according the method prescribed by ^37^. Our measurement at the 25% target was below the *F*_*1/2*_ value of 490 nN reported by Petzold, *et al*., indicating a higher sensitivity when targeting a single position on the animal than when spanning a larger range of the anterior portion of the body.

Figure 5B shows the probability of inducing an acceleration (speed-up) as a function of force and position. Similar to the reversal probability, we found a low probability of inducing a speed-up at the 45% target, further suggesting a gap in sensitivity at the mid-point. Stimuli targeting 75% of the body length caused animals to speed up and the probability of evoking this behavior increased with greater force, similar to the reversal response curve at 25%. We performed the same fit as above to the speed-up probability at 75%. Using the same method of calculating *F*_*1/2*_ as ^25,37^, we found *F*_*1/2*_ = 45.9 nN (95% CI: 32 – 63 nN) for speed-ups at 75%. This metric is an order of magnitude more sensitive than the reported *F*_*1/2*_ for anterior touch ^25^. However, we also observed a lower saturation probability for behaviors evoked by posterior touch than those evoked by anterior touch (Figure 5). In principle, this difference in touch sensitivity could reflect differences in the sensitivity of the anterior and posterior TRNs themselves or in their ability to evoke behavioral responses. In favor of the former possibility, the ALM neurons generate larger mechanoreceptor currents than the PLM neurons ^38^.

### Touch responses are stereotyped motor programs

Once generated, each touch response could fall into one of two behavioral classes: 1) a defined or stereotyped motor response whose quantitative features are uncorrelated to the location or stimulus strength, or 2) a response whose quantitative features depend on stimulus parameters. To distinguish between these possibilities, we analyzed the responses from trials where the response was classified as a reversal or a speed-up across all stimulus strengths. Reversals were taken from trials applied at the stimulus targets of 25% and 35% while speed-ups were taken from stimuli applied at stimulus targets of 75%. From these trials, we analyzed two metrics, 1) *ΔV/V* and 2) maximum acceleration after the stimulus. We determined the distributions of the values for each metric as a function of stimulus strength and target. If the quantitative features of the response were related to the stimulus location and strength, then there should be both a significant difference between the distributions of the metrics and a monotonic relationship between the stimulus strength and the distribution means. If either of those conditions is not met, then the response is likely a stereotyped response.

According to two-way ANOVA, the *ΔV/V* metric is strongly correlated with force (*F*=3.24,*P*=0.007) but not target position (*F* = 1.67, *P* = 0.1964), for the 25% and 35% targets. The two factors did not appear to interact (*F* = 1.35, *P* = 0.2408), and the mean *ΔV/V* did not increase with force (Figure 6A, B). Next, we examined acceleration during each reversal by measuring the maximum backward acceleration of the animal after the stimulus was delivered. Again, considering the 25% and 35% targets, the mean values for this metric were similar at all stimulus strengths and both positions (Figure 6D-E). Two-way ANOVA suggested that stimulus strength and target position affect acceleration (*F* = 2.81, *P* = 0.0163 for stimulus strength; *F* = 8.93, *P* = 0.003 for target position), but the two factors did not appear to interact (*F* = 1.01, *P* = 0.4123). However, mean values for acceleration did not appear to increase or decrease monotonically with force (Figure 6A, B), which suggests that the influence of either force or target position on this metric is weak at best. Since these results do not meet both requirements of statistically significant effects and a monotonic relationship between stimulus strength and distribution means, they indicate that the intensity of reversal has little, if any, dependence on force and is uncorrelated with stimulus position.

**Figure 6.**
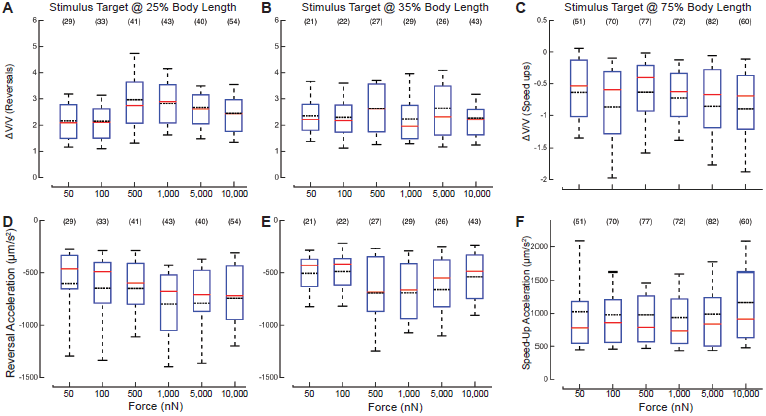
Reversal and speed-up intensity as a function of force suggests that the response is stereotyped motor program. A-C) The distributions of *ΔV/V vs.* force applied for reversals at two stimulus locations, 25% (A), and 35% (B) body length, and for speed-ups at 75% (C) body length. D-F) Distribution of the reversal acceleration *vs.* force at 25% (D), and 35% (E) body length, and forward acceleration at 75% (F) body length. The solid red lines are the median of the distribution; black lines are the mean of the distribution. The bounds of the boxes are the 25% and 75% percentiles. The whiskers are the 10% and 90% percentiles. The sample size is shown in parentheses above each box. A) The distributions of *ΔV/V* according to stimulus strength. B) Distribution of the maximum acceleration during the speed-up according to stimulus strength. The red horizontal line of each box is the median of the distribution. The black dotted line is the mean of the distribution. The bounds of the boxes are the 25% and 75% percentiles. The whiskers are the 10% and 90% percentiles. The sample size is shown above each box.

We also compared the distributions of *ΔV/V* and the maximum forward acceleration during the speed-up according to stimulus strength (Figure 6 C, F). Using a one-way ANOVA to investigate the effect of force, we found no significant effect for either *ΔV/V* or acceleration: *F* = 1.82, *P* = 0.1081 and *F*= 1.07, *P* = 0.3746, respectively. As described above for reversal responses, mean values for accelerations neither increased nor decreased with force (Figure 6 C, F). We conclude that once an animal has decided to reverse or accelerate, it executes a stereotyped motor program whose characteristics are independent of the strength of the stimulus. In other words, the strength of the sensory stimulus governs the probability that a given behavior is executed, but not the nature of the response.

### TRN-specific disruption of spectrin networks impairs touch sensation, but not the touch-evoked motor program

Defects in actin-spectrin networks decrease pre-stress in touch receptor neurons and impair touch sensitivity measured using a classical touch assay ^23^. Such touch impairment is also displayed by *spc-1(dn)* transgenic animals expressing a dominant-negative fragment of *α*-spectrin, disrupting the actin-spectrin network exclusively in the TRNs. We used HAWK to measure the touch defect by delivering stimuli at the 25% target position across a range of stimulus strengths and measuring reversal probability as a function of applied force. We found that the reversal probability for *spc-1(dn)* animals was ∼0.2 for all forces tested (Figure 7A). It remains possible that response probability increases with stronger stimuli. However, this is unlikely since the classical touch assay, which delivers much larger forces than HAWK^13^, revealed a touch defect in these *spc-1(dn)* transgenic animals ^23^.

**Figure 7.**
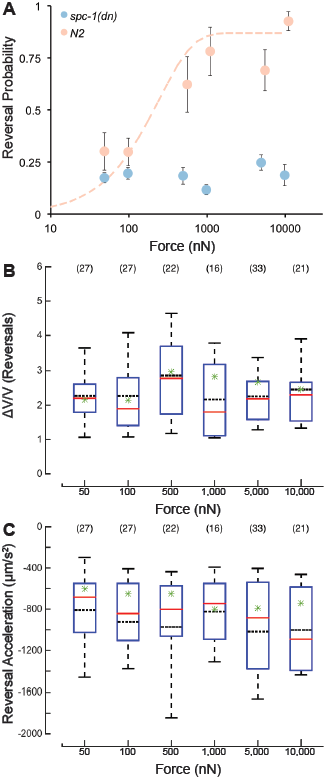
*spc-1(dn)* transgenics have a lower reversal probability across a wide range of force, but reversals that occur are quantitatively similar to wild type. A) Reversal probability *vs.* force of wild type (peach) and *spc-1(dn)* (blue) at 25% body length target. Data for wild type are the same as those in Figure 5A and are re-plotted here for clarity. Points are the mean response rate ± s.e.m. of at least 7 animals at each force-position combination. Each animal was tested during at least 9 trials. B) Distributions of *Δ*V/V *vs.* force. C) Distributions of reversal acceleration *vs.* force. (B-C) Solid, red lines are the median of the distribution; black dotted lines are the mean of the distribution. Green stars indicate the mean for the respective metric from the wild type animal. The bounds of the boxes are the 25% and 75% percentiles. The whiskers are the 10% and 90% percentiles. The sample size is shown in parentheses above each box.

If touch-evoked behaviors are stereotyped, as inferred above, then the non-null trials in which *spc-1(dn)* transgenic animals executed a reversal should have the same quantitative metrics as those found in wild-type animals. Figures 7B and 7C show the distribution of values for the *ΔV/V* metric and reversal acceleration, respectively. As in wild-type animals, neither metric was obviously related to applied force. Two-way ANOVA of the values for the *ΔV/V* metric, with force and genotype as the two factors, revealed a main effect for force (*F* = 3.43, *P*=0.0048), but not for genotype (*F* = 1.93, *P*=0.1657). The two factors did not interact (*F* = 1.21, *P* = 0.303). For the reversal acceleration, the effect of force was not significant (*F* = 1.55, *P* = 0.1729), but the effect of genotype was (*F* = 21.42, *P* = 5.11×10^−6^). Force and genotype did not interact (*F* = 0.69, *P* = 0.6306). In summary, while ANOVA did reveal some statistically significant effects of genotype, there was no monotonic relationship between the mean values and the stimulus strength. These results do not meet both requirements of statistically significant effects and a monotonic relationship between the effect and the metric. Collectively, these results demonstrate that reversals are a stereotyped motor program and that sensory stimulation regulates the probability that the motor program is triggered.

## Discussion

HAWK provides a system for mapping tactile receptive fields in small, freely-moving animals that integrates automated tracking and stimulus targeting, and enables quantitative behavioral analyses. Using animals with known body morphology deviations from wild-type animals, we demonstrated robust targeting by HAWK. Nearly 90% of the trials applied a stimulus within ∼50 μm of the desired target. By integrating a force application system capable of applying forces from 50 nN to 10,000 nN, we used HAWK to perform the first fully quantified, targeted touch assays on freely moving animals. Further, we used the per-frame midline data generated by HAWK to automatically analyze the behavioral response of the animals to stimulus application. Using this analysis we were able to quantitatively map the animals’ touch-sensitive fields and show a stereotyped response independent of a range of stimulus strengths. Although it was designed for application to *C. elegans*, the tracking and stimulus delivery modules could be adapted for application to insect larvae, frog embryos, or leeches. Such an extension would require a refinement of the behavioral analysis module.

Using HAWK, we found that *C. elegans* has anterior and posterior tactile receptive fields for forces < 1,000 nN separated by a gap located near the animal’s mid-point, posterior to the cell body of ALM (Figure 4, Figure 5). This result is consistent with a large body of work showing that *C. elegans* reverses when touched anteriorly and speeds up when touched posteriorly ^4,39^. The existence of a gap between the two receptive fields was unexpected, however. Previous images of the TRNs revealed that the PLM neurite does not overlap with ALM, not even the ALM cell body ^25,40-42^. Given that the gap in sensitivity is between the ALM cell body and the anterior termination of the PLM dendrite, our results suggest that this gap in sensitivity matches the gap in tiling of the skin. We propose that since mechanical strain propagates some distance from the point of stimulation, the gap could allow animals to avoid triggering a reversal when a speed-up is required and *vice versa*. Indeed, mechanical strain propagates tens of microns away from indentation with a 2-4 μm probe and is predicted to propagate further for indentations generated by larger (10 μm) probes ^43^. An error of this kind could be fatal for nematodes seeking to escape a predatory fungus using a lasso-type trap, a behavior known to depend on the TRNs in *C. elegans* ^44^. In this scenario, if the gap is too small, then the nematode risks increasing its exposure to the fungal trap; if the gap is too big, then the nematode may fail to detect trap closure. Thus, there may be a strong evolutionary pressure to optimize the gap size relative to fungal dimensions of ∼10-20 μm^44^.

In addition to a gap in sensitivity, we detected a higher probability of evoking a reversal at the 25% target than at the 35% target (Figure 4, 5). This finding indicates that touch sensitivity is not uniform within a given receptive field and that it is higher near the center than it is near the margins. From prior work, we know that mechanically sensitive ion channels are distributed approximately every 2-4 μm in discrete puncta along neuron processes in *C. elegans* ^25,45-48^. Further, the strain field under an indenter decays to zero at a radius on the order of microns, and is related to the indentation area and indentation depth ^25,43^. Thus, to account for the observed variation in sensitivity within tactile receptive fields, we propose a model in which the probability that an animal will execute a reversal (or speed-up) is proportional to the number of mechanotransduction (MeT) channels activated by a given stimulus. We propose that the number of MeT channels in the strain field produced during touch is a function of both stimulus strength and location. Larger stimuli generate a strain field that could activate more channels. At 25% of body length, there are similar numbers of channels anterior and posterior to the stimulus location available to detect the propagated strain ^25,45-48^. This situation maximizes the number of channels available for activation and may account for higher sensitivity at this position near the center of the receptive field. Targeting 35% of the body length, by contrast, we contact the animal on or proximal to the ALM cell body. However, there are few, if any MeT channels posterior to this position due to the gap in tiling between ALM and PLM ^25,45-48^. As a result, there are fewer channels available for activation, which we propose accounts for our finding of the decreased probability of triggering a response at this position (Figure 5). Finally, targeting 45% of body length results in stimulus delivery posterior to the ALM cell body ^25,45-48^, accounting for the reduced reversal response from the animal in that location.

McClanahan, et al., showed that the pre-stimulus velocity of the animal is uncorrelated to the post-stimulus velocity when initiating a behavior response to a cuticle deflection inside a microfluidic chamber ^14^. We have further demonstrated that the intensities of reversal and speed-up responses, once initiated, are correlated to neither the stimulus strength nor its location (Figure 6), suggesting that stereotyped motor programs govern both reversals and speed-ups. This idea is strengthened by our finding that reversals in a partially touch-insensitive transgenic animal, *spc-1(dn)*, are indistinguishable from those observed in wild-type (Figure 7). Taken together, these findings suggest that the signal delivered by the TRNs regulates the probability that a given motor program will be executed but has no influence on the intensity of the response.

Our results suggest a switch-like model of the mechanosensation pathway: stereotyped behavioral states are activated by an electrochemical switch. Decision making, or transitioning between behavioral states in *C. elegans*, has been described previously using classical switch models, e.g., “single pole/single throw” that starts or stops one circuit/state or “single pole/double throw” that switches between circuits/states ^49^. Faumont modeled the transition from one behavior to another due to a single mechanical stimulus as a single pole/single throw state switch. Our results corroborate this single pole/single model.

TRN signaling is also modulated by strain rate; both direct ion channel recordings and calcium imaging have revealed that MeT current amplitude increases with indentation until it saturates ^36^. Taken together with our results demonstrating stereotyped behavioral responses and the known circuit linking the posterior and anterior TRNs to these motor responses ^6^, we postulate that the interneurons detect the sensory neuron signaling and “switch on” the appropriate behavioral response.

Using calcium imaging, others have determined that specific behavior states such as reversals correspond to the signaling of the specific sets of motor neurons. Whole-brain imaging in *C. elegans* correlates behavior states with interneuron and motor neuron signaling, suggesting that the brain moves between distinct neuronal states correlated with behavior states such as crawling, turning, and reversing^50-52^. Others have reported the direct correlation between motor neuron signaling and behavior states ^53^.Our results are consistent with these imaging and behavior studies in *C. elegans* that concluded that stereotyped behavior states are switched on by upstream neuronal signals.

Other animals used to investigate tactile encoding such as the leech *Hirudo medicinalis* use multiple sensory neurons and neuron types to determine stimulus properties such as location and stimulus strength ^54^. These somatosensory systems rely on interneurons to multiplex and decode signals from multiple classes of sensory neurons. The decoded signal ultimately governs the behavioral response scope; leech responses span global body movements to localized bending ^55,56^. This model aligns with our proposed signaling model in which a set of interneurons is responsible for decoding the sensory neuron signal and triggering the behavioral response.

Using HAWK, this study has established that the mechanosensation pathway behaves as a switch in which the TRNs and interneurons act cooperatively to switch between stereotyped behavior states. Since there are distinct sets of neurons that activate distinct behaviors in *C. elegans*, the mechanosensation pathway consists of two sets of switching circuits: posterior and anterior. Each switch activates the corresponding behavior state: reversal when touched anteriorly or speed-up when touched posteriorly. We establish that *C. elegans* has two tactile receptive fields separated by an insensitive region or sensory gap. Within the anterior receptive field, the threshold for triggering a reversal was lower at the middle of the receptive field (25% of body length) than it was more posteriorly (35%). Thus, sensitivity to mechanical loads is non-uniform within the tactile receptive field covered by the anterior TRNs. Such variations in sensitivity also occur in human tactile receptive fields ^57,58^ and are proposed to be important for tactile object recognition ^59^. From a detailed analysis of the nature and magnitude of behavioral responses, we affirmed that stimulation of the anterior receptive field triggers reversals and that stimulation of the posterior receptive field causes *C. elegans* to speed up. We conclude that both reversals and speed-ups constitute stereotyped motor programs and that analog signals from the TRNs regulate the probability of executing these behaviors. We propose that the gap is essential for nematodes to escape from predatory fungus and its size may be optimized to account for the longitudinal distribution of mechanical strain, ensuring that the two receptive fields function as independent sensors. Our results suggest a switch-like model in which sensory neurons activate distinct stereotyped behavior states, uncorrelated to the strength or location of the stimulus along the sensor neuron. This model distinguishes the roles of sensory and motor neurons into distinct functions and signaling pathways.

## Acknowledgements

The authors thank S. Fechner, M. Krieg, A. Leifer, and C. Fang-Yen for helpful ideation, discussion, and feedback, and Z. Liao for technical support. This work was supported by the National Institutes of Health under Grant R01EB006745 (BLP, MBG), R01NS044715 (MBG), National Science Foundation (NSF) Grant EFRI MIKS 1136790, and the Stanford undergraduate research program. Some *C. elegans* strains were provided by the Caenorhabditis Genetics Center, which is funded by NIH Office of Research Infrastructure Programs (P40 OD010440). Work was performed in part at the Stanford Nanofabrication Facility supported under the NSF NNCI under award ECCS-1542152. There are no conflicts of interest to declare.

## Author contributions

E.A.M., M.G.B., and B.L.P. designed the research, E.A.M, F.L., A.L.N., J.W., B.H., designed and built the system, E.A.M. conducted the experiments, E.A.M., F.L., A.L.N., M.B.G., B.L.P analyzed data and wrote the paper.

## Movies

**Movie 1.**
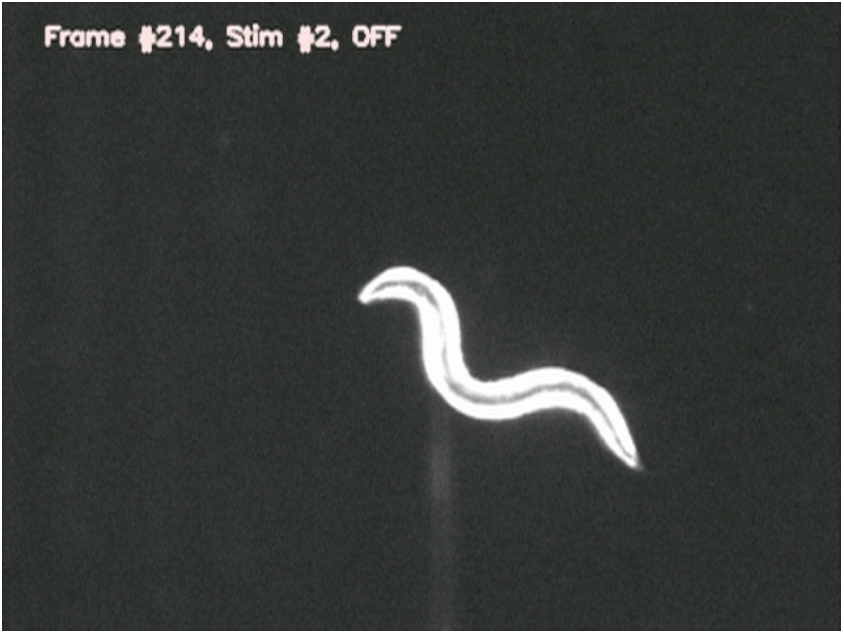
HAWK applies controlled, targeted stimuli to freely moving worms. A series of compiled videos shows four distinct trials delivered on anterior and posterior targets. Frames with the green dot are during the stimulus application. Movie frame rate is 10 fps. Frames are captured at approximately 10 fps, with delays during large stage translations at the beginning and end of stimulus application.

## References

1 S. Cameron and Y. Rao, Mol Brain, 2010, 3, 28.

2 F. Wang, D. P. Julien and A. Sagasti, Cell Adhesion & Migration, 2013, 7, 388–394.

3 M. Petrovic and D. Schmucker, Bioessays, 2015, 37, 996–1004.

4 W. R. Schafer, 2014, 467, 39–48.

5 M. Chalfie, Nat Rev Mol Cell Biol, 2009, 10, 44–52.

6 M. Chalfie, J. E. Sulston, J. G. White, E. Southgate, J. N. Thomson and S. Brenner, Journal of Neuroscience, 1985, 5, 956–964.

7 S. R. Wicks and C. H. Rankin, Journal of Neuroscience, 1995, 15, 2434–2444.

8 A. M. Leifer, C. Fang-Yen, M. Gershow, M. J. Alkema and A. D. T. Samuel, Nat Meth, 2011, 8, 147–152.

9 Y. Cho, D. A. Porto, H. Hwang, L. J. Grundy, W. R. Schafer and H. Lu, Lab Chip, 2017, 17, 2609– 2618.

10 A. L. Nekimken, H. Fehlauer, A. A. Kim, S. N. Manosalvas-Kjono, P. Ladpli, F. Memon, D. Gopisetty, V. Sanchez, M. B. Goodman, B. L. Pruitt and M. Krieg, Lab Chip, 2017, 17, 1116–1127.

11 M. Chatzigeorgiou, L. Grundy, K. S. Kindt, W. H. Lee, M. Driscoll and W. R. Schafer, Journal of Neurophysiology, 2010, 104, 3334–3344.

12 M. Chalfie, A. C. Hart, C. H. Rankin and M. B. Goodman, WormBook, 2014.

13 A. L. Nekimken, E. A. Mazzochette, M. B. Goodman and B. L. Pruitt, 2016, 1–7.

14 P. D. McClanahan, J. H. Xu and C. Fang-Yen, Integr. Biol., 2017, 9, 800–809.

15 S.-J. Park, M. B. Goodman and B. L. Pruitt, PNAS, 2007, 104, 17376.

16 S.-J. Park, B. C. Petzold, M. B. Goodman and B. L. Pruitt, Rev. Sci. Instrum., 2011, 82, 043703.

17 B. C. Petzold, S.-J. Park, P. Ponce, C. Roozeboom, C. Powell, M. B. Goodman and B. L. Pruitt, Biophysical Journal, 2011, 100, 1977–1985.

18 S.-J. Park, J. C. Doll and B. L. Pruitt, J. Microelectromech. Syst., 2010, 19, 137–148.

19 S.-J. Park, J. C. Doll, A. J. Rastegar and B. L. Pruitt, J. Microelectromech. Syst., 2010, 19, 149–161.

20 J. C. Doll, S.-J. Park and B. L. Pruitt, Journal of Applied Physics, 2009, 106, 064310.

21 C. Thacker, J. A. Sheps and A. M. Rose, Cell. Mol. Life Sci., 2006, 63, 1193–1204.

22 T. L. Gumienny, L. T. MacNeil, H. Wang and M. De Bono, Current Biology, 2007, 17, 159–164.

23 M. Krieg, A. R. Dunn and M. B. Goodman, Nature Cell Biology, 2014, 16, 224–233.

24 T. Stiernagle, WormBook, 2006.

25 B. C. Petzold, S.-J. Park, E. A. Mazzochette, M. B. Goodman and B. L. Pruitt, Integr. Biol., 2013, 5, 853–864.

26 J. N. Stirman, M. M. Crane, S. J. Husson, A. Gottschalk and H. Lu, Nat Protoc, 2012, 7, 207–220.

27 A. A. Barlian, W. T. Park, J. R. Mallon, A. J. Rastegar and B. L. Pruitt, Proc. IEEE, 2009, 97, 513– 552.

28 D. R. Brumley, M. Willcox and J. E. Sader, Physics of Fluids (1994-present), 2010, 22, 052001.

29 J. C. Doll and B. L. Pruitt, J. Micromech. Microeng., 2012, 22, 095012.

30 E. A. Mazzochette, 2016.

31 N. Otsu, Systems, Man and Cybernetics, IEEE Transactions on, 1979, 9, 62–66.

32 C. Fang-Yen, M. Wyart, J. Xie, R. Kawai, T. Kodger, S. Chen, Q. Wen and A. D. T. Samuel, PNAS, 2010, 107, 20323–20328.

33 J. Cohen, Educational and Psychological Measurement, 2016, 20, 37–46.

34 G. Cardillo.

35 R. O’Hagan, M. Chalfie and M. B. Goodman, Nat Neurosci, 2004, 8, 43–50.

36 A. L. Eastwood, A. Sanzeni, B. C. Petzold, S.-J. Park, M. Vergassola, B. L. Pruitt and M. B. Goodman, PNAS, 2015, 112, E6955–E6963.

37 C. Mills, D. LeBlond, S. Joshi, C. Zhu and G. Hsieh, The Journal of Pain, 2012, 13, 519–523.

38 X. Chen and M. Chalfie, Journal of Neuroscience, 2015, 35, 2200–2212.

39 M. B. Goodman, WormBook, 2006.

40 M. E. Gallegos and C. I. Bargmann, Neuron, 2004, 44, 239–249.

41 T. Shida, J. G. Cueva, Z. Xu, M. B. Goodman and M. V. Nachury, PNAS, 2010, 107, 21517–21522.

42 D. Lockhead, E. M. Schwarz, R. O’Hagan, S. Bellotti, M. Krieg, M. M. Barr, A. R. Dunn, P. W. Sternberg and M. B. Goodman, Molecular Biology of the Cell, 2016, 27, 3717–3728.

43 M. Elmi, V. M. Pawar, M. Shaw, D. Wong, H. Zhan and M. A. Srinivasan, 2017, 1–12.

44 S. M. Maguire, C. M. Clark, J. Nunnari, J. K. Pirri and M. J. Alkema, Current Biology, 2011, 21, 1326–1330.

45 L. Emtage, G. Gu, E. Hartwieg and M. Chalfie, Neuron, 2004, 44, 795–807.

46 J. G. Cueva, A. Mulholland and M. B. Goodman, Journal of Neuroscience, 2007, 27, 14089–14098.

47 J. Arnadottir and M. Chalfie, Annu. Rev. Biophys., 2010, 39, 111–137.

48 V. Vásquez, M. Krieg, D. Lockhead and M. B. Goodman, Cell Reports, 2014, 6, 70–80.

49 S. Faumont, T. H. Lindsay and S. R. Lockery, Current Opinion in Neurobiology, 2012, 22, 580– 591.

50 S. Kato, H. S. Kaplan, T. Schrödel, S. Skora, T. H. Lindsay, E. I. Yemini, S. R. Lockery and M. Zimmer, Cell, 2015, 163, 656–669.

51 J. P. Nguyen, F. B. Shipley, A. N. Linder, G. S. Plummer, M. Liu, S. U. Setru, J. W. Shaevitz and A. M. Leifer, PNAS, 2016, 113, E1074–E1081.

52 V. Venkatachalam, N. Ji, X. Wang, C. Clark, J. K. Mitchell, M. Klein, C. J. Tabone, J. Florman, H. Ji, J. Greenwood, A. D. Chisholm, J. Srinivasan, M. J. Alkema, M. Zhen and A. D. T. Samuel, PNAS, 2016, 113, E1082–8.

53 V. J. Butler, R. Branicky, E. I. Yemini, J. F. Liewald, A. Gottschalk, R. A. Kerr, D. B. Chklovskii and W. R. Schafer, Journal of The Royal Society Interface, 2014, 12, 20140963.

54 F. Pirschel and J. Kretzberg, Journal of Neuroscience, 2016, 36, 3636–3647.

55 S. M. Baca, E. E. Thomson and W. B. Kristan Jr., Journal of Neurophysiology, 2005, 93, 3560– 3572.

56 E. E. Thomson and W. B. Kristan, J. Neurosci., 2006, 26, 8009–8016.

57 R. S. Johansson, J. Physiol. (Lond.), 1978, 281, 101–125.

58 J. R. Phillips, R. S. Johansson and K. O. Johnson, Journal of Neuroscience, 1992, 12, 827–839.

59 J. A. Pruszynski and R. S. Johansson, Nat Neurosci, 2014, 17, 1404–1409.

